# Excavating New Facts from Ancient Hepatitis B Virus Sequences

**DOI:** 10.1101/829473

**Authors:** Sibnarayan Datta

## Abstract

Recently, Muhlemann et al. (2018) and Krause-Kyora et al. (2018) discovered 15 ancient Hepatitis B virus (aHBV) sequences dating back to the Neolithic age (NA) and the Bronze age (BA). Being published simultaneously, neither of these studies could include sequences from the other for analyses. In the present research, aHBV sequences from these studies were collective re-analysed with reference to a comprehensive database comprising extant HBV diversity to understand their relatedness and role in the evolution of extant HBV diversity. Present analyses revealed several interesting findings on distribution, dispersal, phylogenetic and recombinational relatedness of ancient HBV to extant genotypes, which were not recognized previously. Several interesting recombination patterns were observed, which corroborated well with ancient human migration, shown by the human genetic studies. Present analyses suggest that comparable to the replacement of the Neolithic European farmer associated Y chromosome haplogroups by haplogroups associated with the steppe people during *Steppe migration*, HBV genotype associated with the early Neolithic European farming cultures was also replaced by the ancestral HBV genotype A probably carried by the migrating steppe people, and a variant of this genotype is the prevalent HBV genotype in contemporary European populations. Additionally, based on recent literature, this research also indicates that HBV genotype divergence estimates proposed by Muhlemann et al., and others cannot sufficiently explain distribution of certain extant HBV genotypes. Hence, an alternative possibility to explain long distance and trans-oceanic distribution of phylogenetically related HBV genotypes was reviewed and discussed in the light of currently available knowledge. Through this manuscript, novel and important findings of the present analyses are communicated.

---

Rapid technological advancements in sequencing technologies and computational tools over the last few decades have enabled sequencing and reconstruction of highly divergent genomes including ancient pathogen genomes (*Spyrou et al., 2019*). This has established the field of ‘*ancient pathogen genomics’*, which is helping us better understand their origin, evolution, epidemiology apart from revealing their interaction with hosts (*Spyrou et al., 2019*). Recently, *Mühlemann and colleagues* reported 12 complete or partial ancient Hepatitis B virus (aHBV) genome sequences isolated from archaeological human remains dating back to ca. 4500 years. Concurrently, *Krause-Kyora and colleagues*, reported three additional aHBV sequences, two of which were more ancient (ca. 5000-7000 years) than *Mühlemann et al*. For the first time, these two studies provided direct evidence for human HBV infection since Neolithic and Bronze ages. However, being published simultaneously, neither of these studies could include sequences from the other for comparative analyses. Additionally, two other previous studies provided aHBV from recent past (ca. 400-500 years) (*Pattersson et al., 2018; Bar-gal et al., 2012*). These sequences are precious, as they offer a unique opportunity to study the evolution and distribution of HBV through ancient ages and to better understand the distribution of extant HBV diversity. In the research presented in this manuscript, available aHBV sequences were re-analysed collectively, results were interpreted, reviewed and discussed in the light of available knowledge.

aHBV and extant HBV sequence used in this analysis were either received from the original authors or retrieved from the NCBI GenBank. Sequences were organized using BioEdit v7 (*Hall, 1999*). To ensure that the working dataset represent extant genotypes/subgenotypes in a proportionate and phylogenetically more informative manner in molecular evolutionary analyses, identical and overrepresented sequences (particularly human HBV genotypes A-D) were removed cautiously. The final dataset comprised of 17 aHBV DNA sequences (Table 1) reported till date (*Spyrou et al., 2019; Mühlemann et al., 2018; Krause-Kyora et al., 2018; Pattersson et al., 2018; Bar-gal et al., 2012*), along with 258 well characterized sequences representing extant human genotypes/subgenotypes classified till date and 43 sequences representing non-human primate (NHP) associated HBV (NHP-HBV) sequences (*Kramvis, 2014*). Sequence dataset was aligned using the MAFFT algorithm (*https://mafft.cbrc.jp/alignment/server/*). Best fitting nucleotide substitution model (GTR+G+I) was determined based on the Bayesian Information Criterion (BIC), the corrected Akaike Information Criterion (AIC), and maximum likelihood parameters, using MEGA v6 (*Tamura et al., 2013*). Phylogenetic network and phylogenetic tree were reconstructed using SplitsTree v4 (*Huson & Bryant, 2006*) and MEGA, respectively. Reliability of the phylogenetic network and the tree were verified by bootstrap analyses (at least 1000 iterations). In contrast to the previous study (*Mühlemann et al., 2018*), in this analysis, phylogenetic network model was preferred over simple bifurcating tree model, as networks enable effective presentation of complicated evolutionary scenarios, taking into account reticulate events such as horizontal gene transfer, hybridization, recombination etc. that are expected to be involved (*Huson & Bryant, 2006; Makarenkov et al., 2004*). Recombination was examined using different algorithms implemented in RDP v4 program (*Martin et al., 2015*). Recombination events supported by an average P value < 0.01 and detected by at least three different algorithms were considered true events.

**Table 1.**
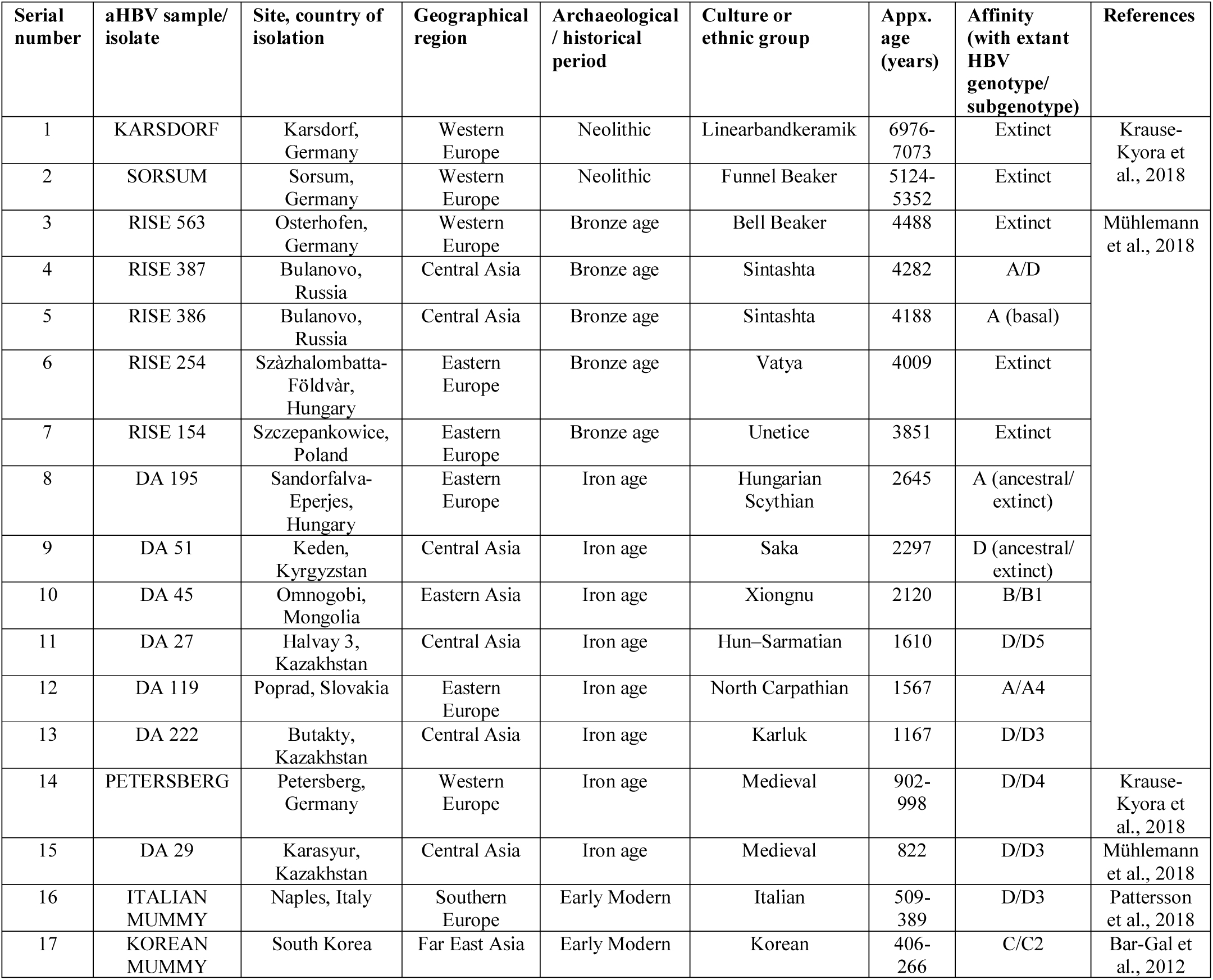
Details of ancient HBV (aHBV) sequences reported till date (arranged chronologically).

| Serial number | aHBV sample/ isolate | Site, country of isolation | Geographical region | Archaeological / historical period | Culture or ethnic group | Appx. age (years) | Affinity (with extant HBV genotype/ subgenotype) | References |
| --- | --- | --- | --- | --- | --- | --- | --- | --- |
| 1 | KARSDORF | Karsdorf, Germany | Western Europe | Neolithic | Linearbandkeramik | 6976-7073 | Extinct | Krause-Kyora et al., 2018 |
| 2 | SORSUM | Sorsum, Germany | Western Europe | Neolithic | Funnel Beaker | 5124-5352 | Extinct |  |
| 3 | RISE 563 | Osterhofen, Germany | Western Europe | Bronze age | Bell Beaker | 4488 | Extinct | Mühlemann et al., 2018 |
| 4 | RISE 387 | Bulanovo, Russia | Central Asia | Bronze age | Sintashta | 4282 | A/D |  |
| 5 | RISE 386 | Bulanovo, Russia | Central Asia | Bronze age | Sintashta | 4188 | A (basal) |  |
| 6 | RISE 254 | Százhalombatta-Földvár, Hungary | Eastern Europe | Bronze age | Vatya | 4009 | Extinct |  |
| 7 | RISE 154 | Szczepankowice, Poland | Eastern Europe | Bronze age | Unetice | 3851 | Extinct |  |
| 8 | DA 195 | Sandorfalva-Eperjes, Hungary | Eastern Europe | Iron age | Hungarian Scythian | 2645 | A (ancestral/ extinct) |  |
| 9 | DA 51 | Keden, Kyrgyzstan | Central Asia | Iron age | Saka | 2297 | D (ancestral/ extinct) |  |
| 10 | DA 45 | Omnogobi, Mongolia | Eastern Asia | Iron age | Xiongnu | 2120 | B/B1 |  |
| 11 | DA 27 | Halvay 3, Kazakhstan | Central Asia | Iron age | Hun–Sarmatian | 1610 | D/D5 |  |
| 12 | DA 119 | Poprad, Slovakia | Eastern Europe | Iron age | North Carpathian | 1567 | A/A4 |  |
| 13 | DA 222 | Butakty, Kazakhstan | Central Asia | Iron age | Karluk | 1167 | D/D3 |  |
| 14 | PETERSBERG | Petersberg, Germany | Western Europe | Iron age | Medieval | 902-998 | D/D4 | Krause-Kyora et al., 2018 |
| 15 | DA 29 | Karasyur, Kazakhstan | Central Asia | Iron age | Medieval | 822 | D/D3 | Mühlemann et al., 2018 |
| 16 | ITALIAN MUMMY | Naples, Italy | Southern Europe | Early Modern | Italian | 509-389 | D/D3 | Pattersson et al., 2018 |
| 17 | KOREAN MUMMY | South Korea | Far East Asia | Early Modern | Korean | 406-266 | C/C2 | Bar-Gal et al., 2012 |

In the phylogenetic network (NeighborNet) analysis (Figure 1), two Neolithic sequences (KARSDORF, SORSUM) and three Bronze age aHBV sequences (RISE563, RISE254, RISE154), together represented a clade of an human HBV genotype (presently extinct), that arose from the main stem of African clade of NHP-HBV, agreeing to the original studies. Within this clade, two distinct but deeply separated clusters were visible further – (i) KARSDORF + RISE154 and (ii) SORSUM + RISE563 + RISE254. On the other hand, RISE387 occupied an intermediate position between genotype A and D clades, while the basal positions of RISE386 and DA51 in the genotype A and D clades, respectively, were suggestive of these sequences representing presently extinct subgenotypes or ancient ancestral sequence for their corresponding genotypes (Figure 1). This finding appears to be consistent with the proposed origin for genotype A in central Asia (*Kostaki et al., 2018*) from where RISE386 was isolated (Bulanovo, Russia), as suggested earlier (*Muhlemann et al., 2018*). Genetic affinities of remaining aHBV sequences included in this analysis agree with the original reports (Figure 1, Figure 2). Nevertheless, widespread reticulation was evident at the base of the phylogenetic network, which reduced drastically once specific genotype clades began diverging (Figure 1), indicating extensive sequence swapping among primordial HBV sequence that subsequently led to emergence and evolution of extant genotypes. Furthermore, absence of any recombination ‘*bottleneck*’ in the phylogenetic network signified a relatively unbiased geographical distribution and expansion of the aHBV genomes. Therefore, present analysis reaffirms previous findings that majority of the extant HBV genotypes are recombinants, many of which were conventionally regarded as ‘*non-recombinant*’ (*Simmonds & Midgley, 2005*).

**Figure 1.**
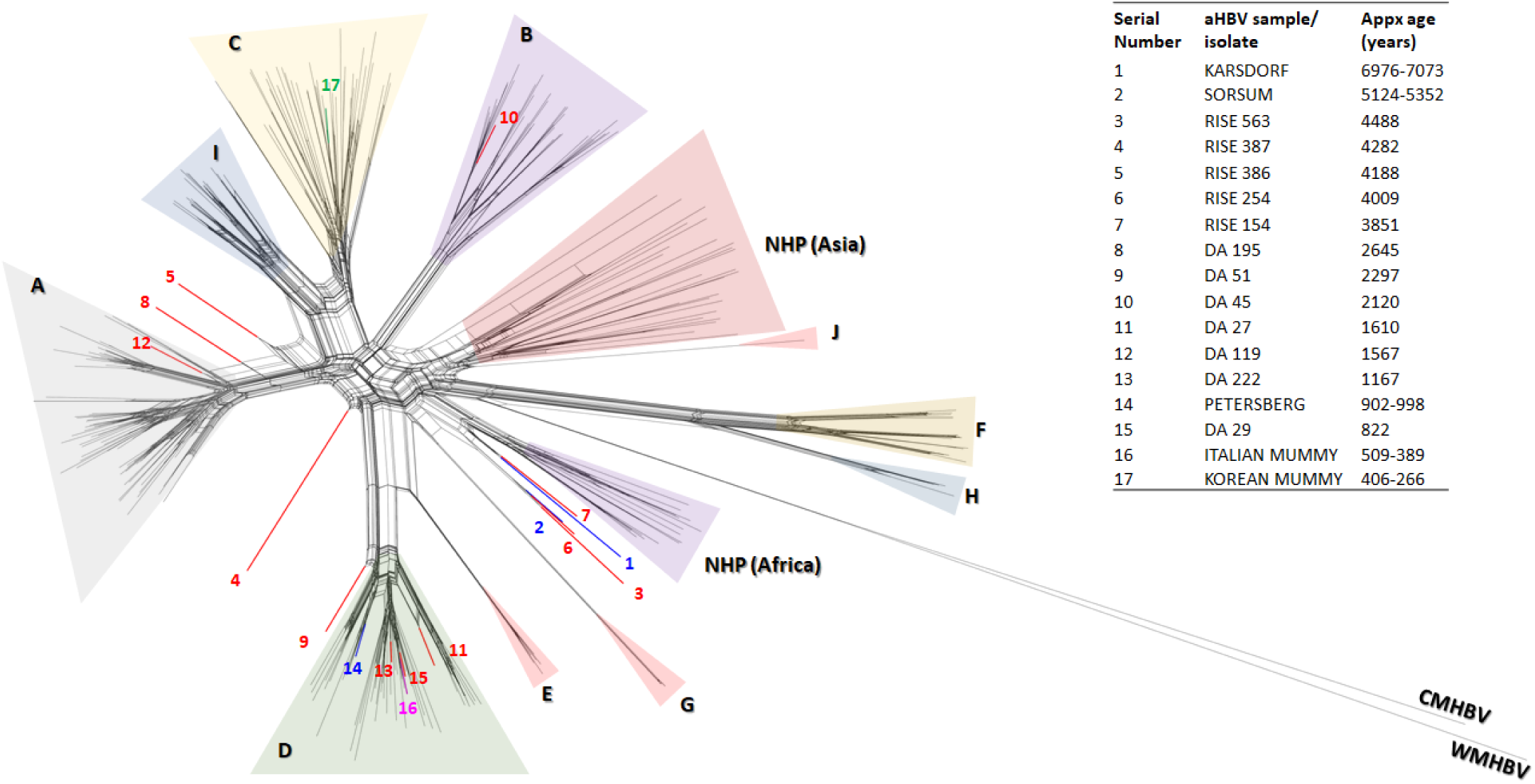
NeighborNet phylogenetic network showing relatedness among 17 aHBV, 258 extant human HBV and 43 non-human primate (NHP) complete genome sequence. Clades overlaid by different colours indicate ten currently classified human HBV genotypes (A–J), two NHP HBV genotypes (African and Asian). aHBV sequences are represented by coloured branches (same colours indicate sequences from same study). Number of the coloured branches correspond to the serial number of aHBV sequences listed in the adjacent table. CMHBV, capuchin monkey HBV; WMHBV, woolly monkey HBV.

**Figure 2.**
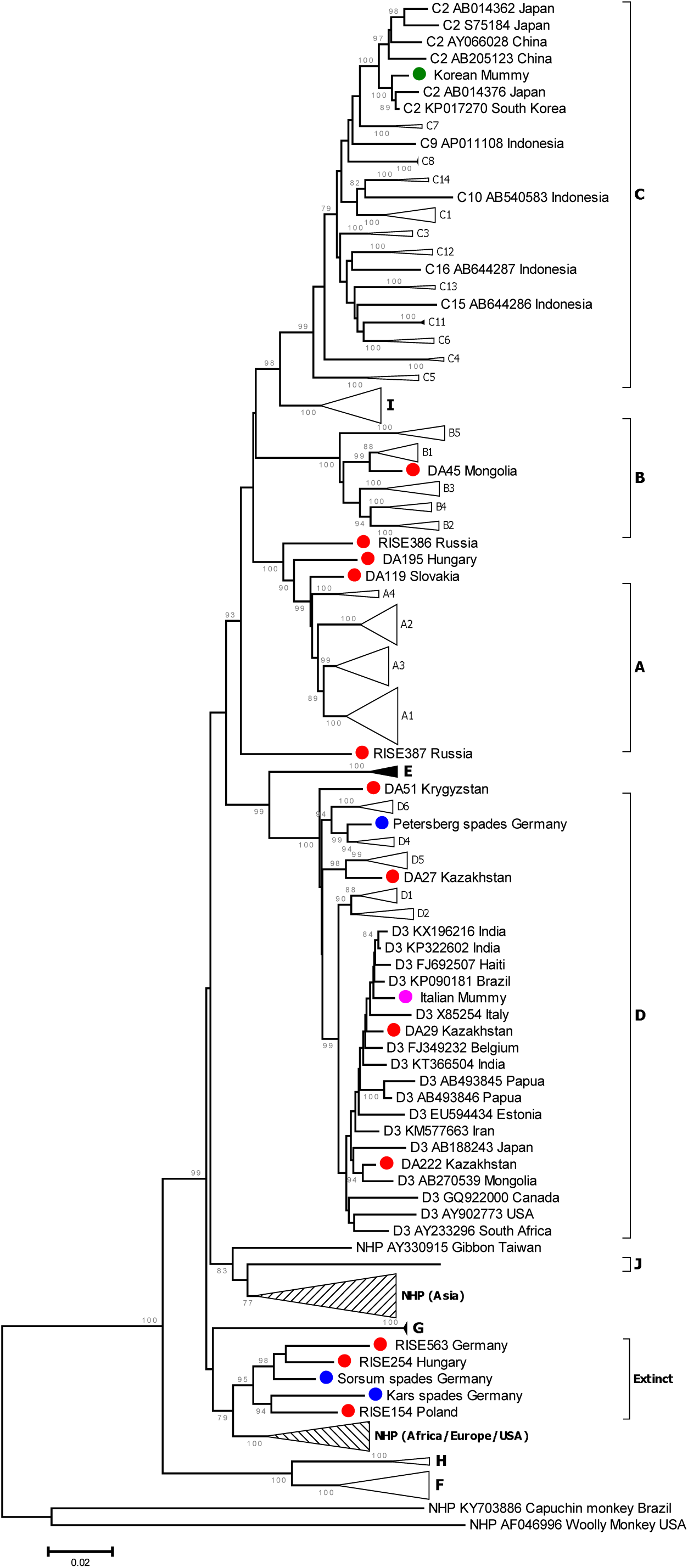
Neighbour-Joining tree representing phylogenetic relatedness among aHBV and extant HBV genotypes/ subgenotypes. Branch lengths represent evolutionary distances, computed using the Maximum Composite Likelihood (MCL) method. The percentage of replicate trees in which the related taxa clustered together during the bootstrap test (1000 replicates) are shown at branch nodes. aHBV sequences are represented by solid coloured spheres (same colours indicate sequences from same study). For better representation of the relevant information, some branches in the tree, which are not pertinent in the current perspective, have been collapsed.

To complement the previous findings (*Muhlemann et al., 2018*), analysis of recombination in the aHBV sequences was done with reference to a considerably large dataset (mentioned in previous section) rationally organised to represent the existing global HBV diversity in a proportionate manner. Recombination breakpoints and parental sequences identified in several HBV sequences included in the present analysis matched with previous analyses (*Muhlemann et al., 2018; Simmonds & Midgley, 2005*), authenticating the results of present analyses. Since recombination detection methods depend upon the presence of probable parental sequences in the reference dataset, inclusion of available aHBV sequences in the present dataset helped identify several events, that could not be identified in previously studies. The present analyses identified a SORSUM like sequence to be the major contributor in ancient and existing genotype A sequences (Table 2, Figure 3), while a RISE563 like sequence in all the existing genotype I sequences (Table 3), both of which could not be unidentified in the previous studies (*Muhlemann et al., 2018; Simmonds & Midgley, 2005*). However, considering the current availability of a limited number of aHBV sequences, it was assumed here that event showing involvement of temporally widespread parental sequences denote the involvement of yet unknown phylogenetically closely related ancestors or descendants of these aHBV sequences, therefore the word ‘like’ is used hereafter to describe such events. Nevertheless, detection of sequences similar to African NHP HBV as major contributors in Neolithic and early Bronze age aHBV sequences, with minor contributions from sequences similar to extant human HBV genotypes (events RA, RD in Table 2, Figure 3, Figure 4), appear to imply African origin of HBV and exchange of HBV sequences among the ancestral human and primates, supporting previous suppositions (*Littlejohn, 2016; Kramvis et al., 2014; Locarnini, 2013*). Recombination fragments like genotype D (subgenotype D6) and genotype E (Table 3, Figure 4) in the aHBV sequences is also apprehensible, considering the prevalence of these genotypes/subgenotypes in Africa (*Littlejohn, 2016; Kramvis et al., 2014; Locarnini, 2013*). On the contrary, detection of fragments like genotype C (subgenotype C2) and gibbon HBV sequences (both being presently confined to far-east and south East Asia respectively), in aHBV isolates from Neolithic and Bronze Age cultures of Europe/central Asia was rather unexpected (events RA, RD in Table 2, Figure 3, Figure 4).

**Table 2.**
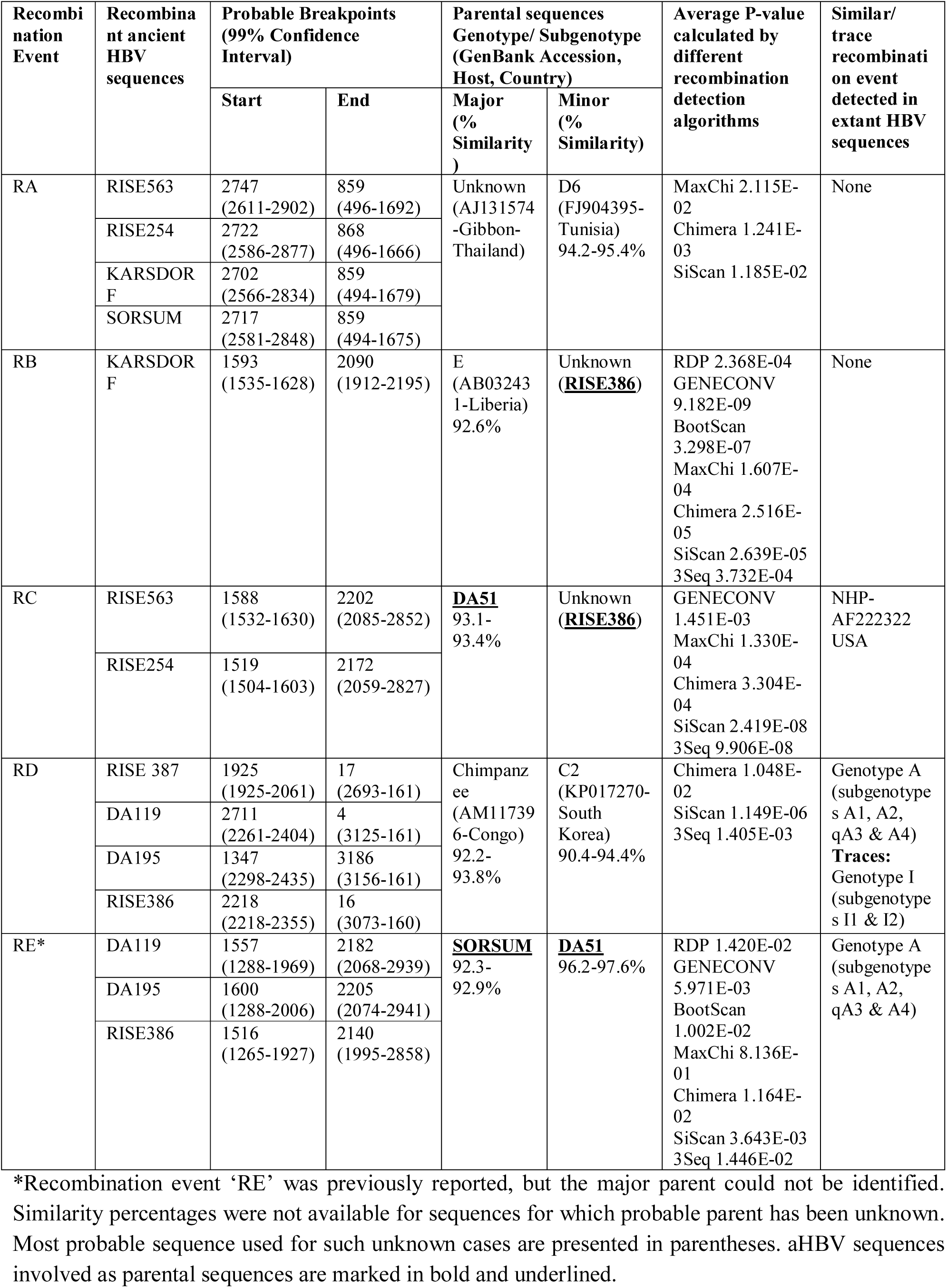
Details of aHBV recombination events detected in the present analysis.

**Table 3.** Details of recombination events involving aHBV sequences as one of the possible parental sequence, observed in extant sequences.

| Extant Recombinant sequence (s) | Probable Breakpoints (99% Confidence Interval) |  | Parental sequences Genotype/ Subgenotype (GenBank Accession, Host, Country) |  | Average P-value calculated by different recombination detection algorithms |
| --- | --- | --- | --- | --- | --- |
|  | Start | End | Major (% similarity) | Minor (% similarity) |  |
| Genotype E; D3 (AB270539-Mongolia); D3 (AY233296-South Africa); D3 (AY902773-USA); D2 (JF754597-Turkey); **Partial evidence:** C2 (KP017270-South Korea) **Trace evidence:** C5 (AF241410-Vietnam); AB241110-Philippines; C14 (HM011493-Indonesia); C3 (X75656-Polynesia & AF458664-China); C11; C15) | 1738-2150 | Undetermined | **<u>SORSUM</u>** 92.9% | D4 (FJ692536-Haiti) 96.7-98.4% | GENECONV 9.235E-03<br>BootScan 2.611E-03<br>MaxChi 6.306E-02<br>Chimera 4.488E-03<br>SiScan 1.289E-12<br>3Seq 1.254E-02 |
| Genotype E (Exclusive) | 2282-2545 | 3166-543 | **<u>RISE386</u>** 92.5% | C6 (AB493842-Indonesia) | MaxChi 4.126E-03<br>Chimera 8.460E-03<br>SiScan 1.246E-12<br>3Seq 2.123E-02 |
| Genotype I (Exclusive) | 1425-1673 | 2898-6 | **<u>RISE563</u>** 92.6% | C2 (KP017270-South Korea) 95% | GENECONV 2.072E-06<br>Bootscan 5.093E-03<br>MaxChi 2.049E-06<br>Chimera 5.755E-06<br>Siscan 2.454E-18<br>3Seq 1.716E-04 |
| Genotype G (Exclusive) | 620-1625 | 1897-2898 | E-X75664-Senegal | Unknown (**<u>RISE386</u>**) | GENECONV 6.937E-11<br>MaxChi 2.870E-05<br>Chimera 2.636E-03<br>SiScan 1.020E-09 |
| NHP-AF498266 (Chimpanzee-Africa) | 492-618 | 958-1061 | NHP (AM117394-Chimpanzee-Cameroon) 95.8% | C2 (AB014376-Japan) 98.3% | RDP 1.171E-08<br>GENECONV 8.449E-10<br>BootScan 6.777E-09<br>MaxChi 1.349E-02<br>Chimera 3.167E-03<br>SiScan 1.804E-05<br>3Seq 4.283E-10 |
| NHP-AF222322 (Chimpanzee-USA) | 1540-1639 | 2095-2863 | **<u>DA51</u>** 90.6% | Unknown (**<u>RISE386</u>**) | GENECONV 1.451E-03<br>MAxChi 1.330E-04<br>Chimera 3.304E-04<br>SiScan 2.419E-08<br>3Seq 9.906E-08 |
| C4 (AB048704 & AB048705)- Australia | 3101-299 | 970-1248 | C2 (AB014376-Japan) 94.7% | NHP (AY781180-Gibbon Cambodia) 92.2% | MaxChi 4.640E-05<br>Chimera 2.030E-02 |

Similarity percentages were not available for sequences for which probable parent(s) were unknown. Most probable sequence used for such unknown cases are presented in parentheses. aHBV sequences involved as parental sequences are marked in bold and underlined.

**Figure 3.**
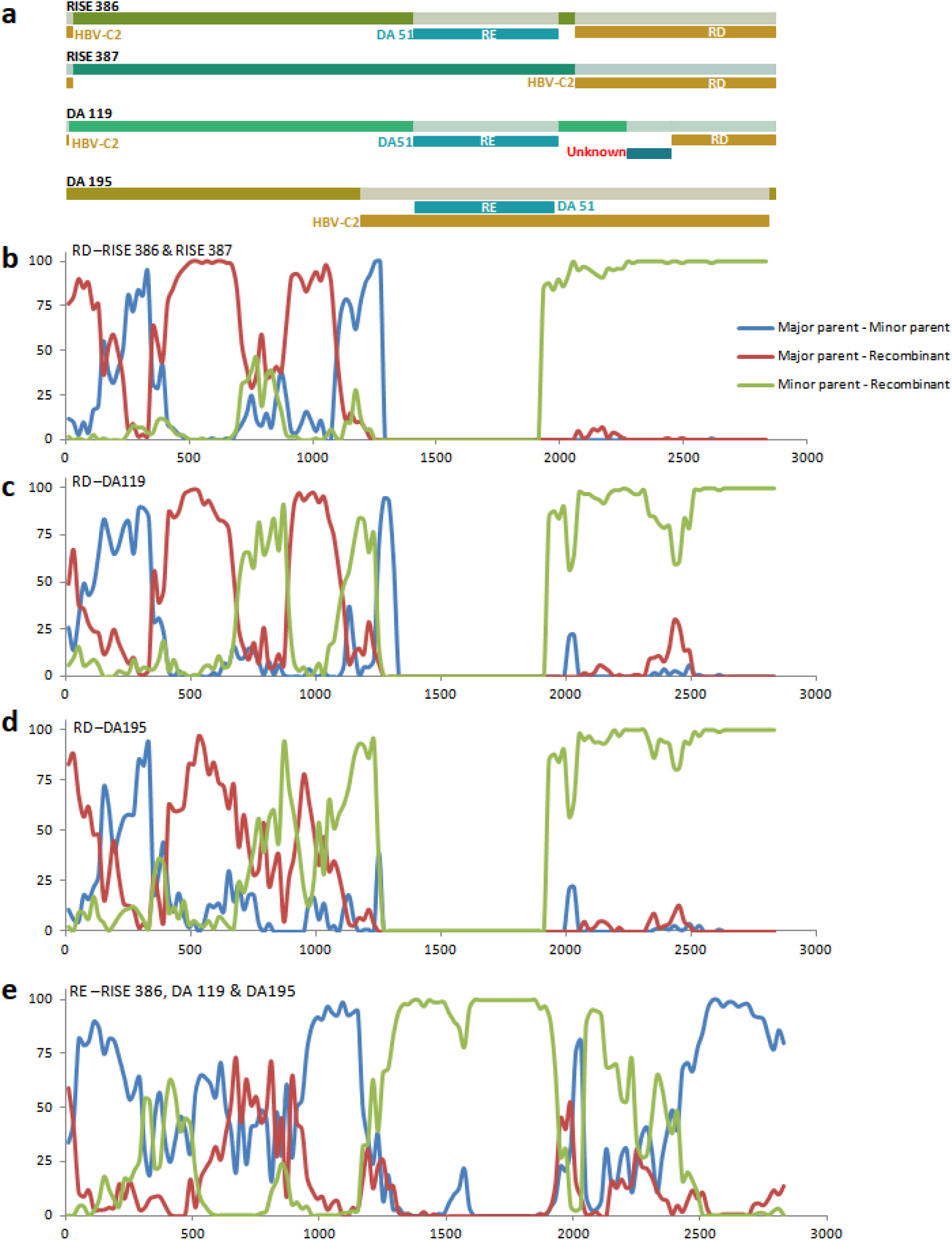
Detailed analyses of recombination events detected in the ancient aHBV sequences, isolated from Bronze and Iron Age (belonging to the genotype A). **a**, schematic representation of the recombination break points. Recombination events are denoted by RD and RE, details of which are provided in the Table 2. **b**, Bootscan plot of the event RD in isolates RISE386 & RISE387. **c**, Bootscan plot of the event RD in isolate DA119. **d**, Bootscan plot of the event RD in isolate DA195. **e**, Bootscan plot of the event RE in isolates RISE386, DA119 & DA195. In the Bootscan plots, *y* axis represents bootstrap support (%) and *x* axis represent nucleotide position in the HBV genome. Major and minor parents involved in each of these events are detailed in the Table 2.

**Figure 4.**
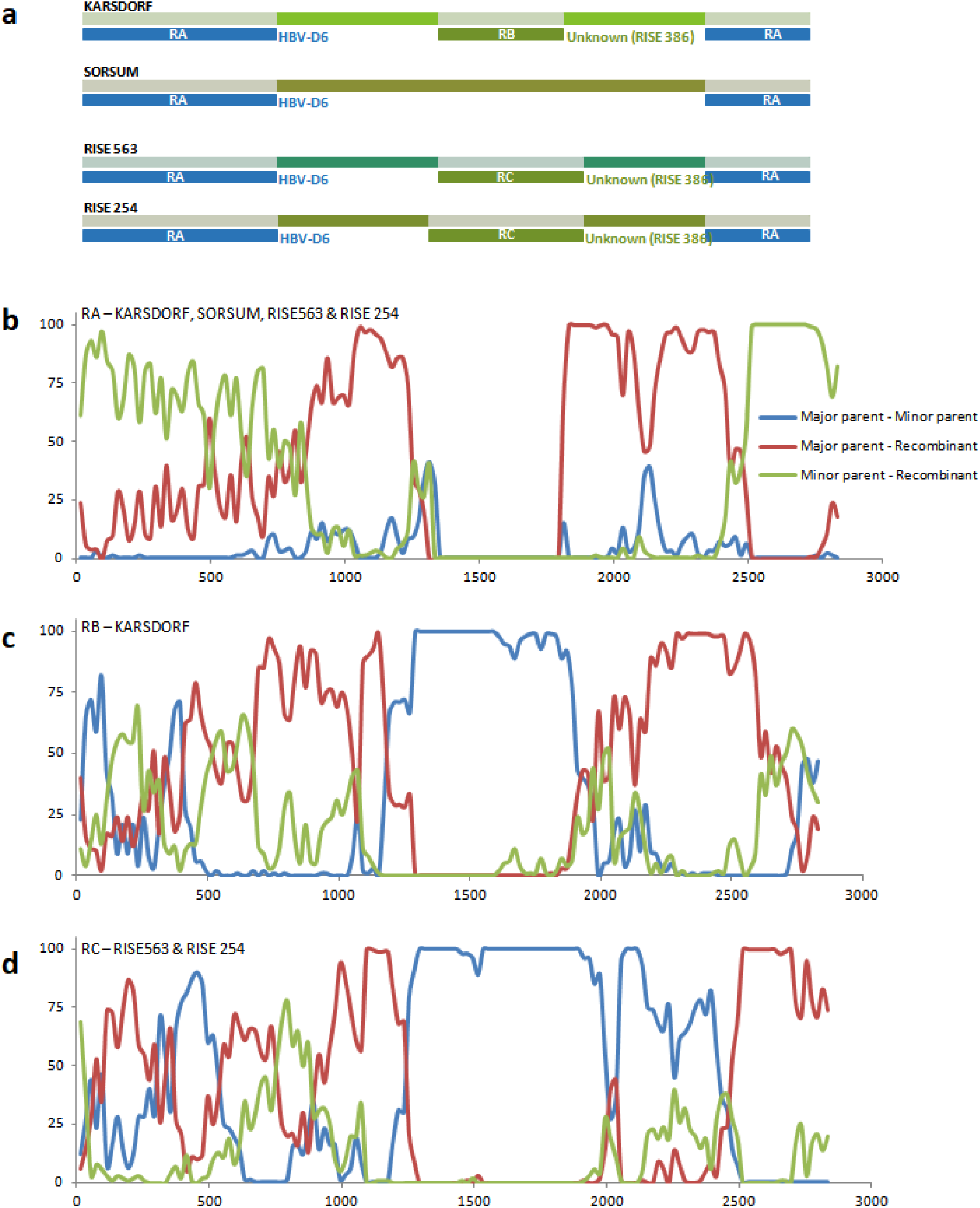
Detailed analyses of recombination events detected in the ancient aHBV sequences, isolated from Neolithic and Bronze Age (belonging to the extinct genotype). **a**, schematic representation of the recombination break points. Recombination events are denoted by RA, RB and RC, details of which are provided in the Table 2. Most probable sequence used for unknown cases are presented in parentheses. **b**, Bootscan plot of the event RA. **c**, Bootscan plot of the event RB. **d**, Bootscan plot of the event RC. In the Bootscan plots, *y* axis represents bootstrap support (%) and *x* axis represent nucleotide position in the HBV genome. Major and minor parents involved in each of these events are detailed in the Table 2. HBV-D6 denotes sequence similar to subgenotype D6 of genotype D.

While searching for similar recombination in the currently available literature, recombination events involving subgenotype C2 sequence was found in at least one wild caught east African chimpanzee (isolate FG, GenBank accession No. AF498266), which has been suggested to indicate genotype C to be an ancient genotype that was hitherto present in Africa (*Littlejohn, 2016; Magiorkinis et al., 2005; Vartanian et al. 2002*). The ancient origin of genotype C is also supported by the observations that this genotype flaunts highest genetic diversity among all other extant human HBV genotypes (16 distinct subgenotypes classified till date) and its recombination with majority of the human and nonhuman primate genotypes predominant in distant geographical regions (*Kramvis, 2014; Magiorkinis et al., 2005; Simmonds and Midgley, 2005*). Furthermore, diverse subgenotypes of this genotype are detected in certain ‘*relict*’ hunter-gatherer populations that are believed to be the descendants of indigenous east African people, which populated different islands of south east Asia and Australia, more than 50 thousand years ago (*Bellwood et al., 2018; Littlejohn, 2016; Kramvis, 2014; Locarnini et al., 2013*). The present findings from aHBV analysis showing subgenotype C2 recombinant sequence in a genetic region discrete from that previously reported in chimpanzee FG (*Magiorkinis et al., 2005*), also support the prehistoric circulation of genotype C genomes (probably intact) in Africa and that further implies that its interaction with chimpanzee HBV took place at least more than once. It is noteworthy to mention that in a recent analyses subgenotype C2 (now termed as quasi-subgenotype C2) has been found to be a non-monophyly (*Kramvis, 2014; Pourkarim et al., 2014; Shi et al., 2012*), which makes the origin of this subgenotype even more elusive. Similarly, presence of gibbon HBV like sequence (presently predominant in south east Asia) in Neolithic aHBV from Europe might be a resultant of ancient interspecies HBV transmissions. Similar interspecies transmission or presence of human HBV genotypes in NHPs that do not correspond to their present-day geographic distributions have also been documented previously (*Rasche et al., 2016; Simmonds, 2001*). This is apprehensible from the recent estimates that primeval HBV evolved several million years ago (mya, probably in Africa, long before the divergence and global dispersal of primate/ human ancestors (*Dominguez Souza et al., 2018; Lauber et al., 2017*). Though the global dispersal of human genotypes is consistent with the conventional ‘*Out of Africa*’ model, dispersal of primate genotypes, is still a matter of hot debate, since the existing models necessitate the ancestors of these primates to undertake improbably long distance trans-Atlantic or intercontinental voyages.

In this analysis, most of the aHBV sequences showed evidences of genetic exchange with RISE386 and/or DA51 like sequences, the two being ancestral representatives for genotype A and D, respectively (events RB, RC, RE in Table 2). Based on their findings, *Mühlemann et. al.* suggested that an ancestor of DA51 recombined with an ‘*unknown*’ major parent to form genotype A. In the present analysis, SORSUM like aHBV was identified as the ‘unknown’ major parent (event ‘RE’, Table 2, Figure 3), and the SORSUM/DA51 like recombinant sequence was evident in all the extinct (DA119, DA195, RISE386) and extant subgenotypes of genotype A (A1, A2, qA3, A4 sequences included in the dataset), signifying that the extant subgenotypes of A emerged subsequent to the event ‘RE’. The recombination breakpoints determined in the present analyses closely corresponded to that inferred in the original study (*Mühlemann et al*., *2018*). However, contrasting to *Mühlemann et al*., event RE could not be recognized in isolate RISE387 in the present analysis, possibly due to a data gap of ∼45 bases, immediately adjacent to the predicted starting breakpoint. It is worth notice that both the events RD & RE were detected in all the subgenotypes of A, while traces of only event RD was present in most of the extant genotype I subgenotypes (I1, I2) included in the present dataset. This similarity of recombination fragment along with phylogenetic network suggests that recombination event RD took place before the divergence of common ancestor of genotypes A and I, while event RE later took place exclusively in the genotype A lineage, supporting previous findings (*Mühlemann et al., 2018*). This piece of direct evidence substantiates earlier propositions of ancient origin of genotype I, which were primarily based on the observed restricted distribution pattern of this genotype in diverse geographical regions across SE Asia (Vietnam, Laos, southern China) and in certain primitive tribal populations of India having no known epidemiological contact with the outside world (*Haldipur et al., 2014; Arankalle et al., 2010; Fang et al., 2011; Su et al., 2014; Olinger et al., 2008; Tran et al., 2008; Hannoun et al., 2000*). Previously, genotype I was proposed to represent link between European/African and Asian HBV genotypes that originated outside South East Asia through complex ancient intergenotypic recombination between genotype C and ‘an *unknown genotype* or a *divergent* or *extinct subgroup of genotype A*’ (*Araujo et al., 2015; Olinger et al., 2008; Hannoun et al., 2000*). The present recombination analysis of genotype I revealing extinct RISE563 like sequence as the major contributor, subgenotype C2 and recombinant fragments like ancestral genotype A aptly agree to the earlier proposal. Additionally, evidences of aHBV sequences as a contributing parent in extant HBV genotype/subgenotypes were also documented in the present analysis (Table 3).

Collective re-analysis of aHBV sequences disclose other important facts about HBV diversity and distribution in ancient Eurasia, which nicely corroborates with the history of human migrations/ movements and invasions. From the available data, it is apparent that the presently extinct Neolithic HBV genotype (represented by KARSDORF, SORSUM, RISE563, RISE254, RISE154) was distributed in western and eastern Europe (present day Germany, Poland and Hungary) for at least 3000 years (7000-4000 YBP) spanning Neolithic cultures (in the early Neolithic farming cultures - Linearbandkeramik and Funnel-Beaker) to Bronze age European cultures (Bell-beaker, Vatya and Unetice). Parallelly, during the Bronze age, a distinct HBV genotype (represented by RISE386, an ancestral genotype related to present day genotype A) appeared in the Sintashta culture from the Eurasian steppe, bordering Eastern Europe and Central Asia. Further, RISE386 from the Sintashta culture showed recombination between Neolithic sequence (SORSUM like major parent) and an ancestral HBV sequence related to extant genotype D (DA51 like minor parent). The proposed origin of Sintashta people from eastward migration of the Corded Ware people (believed to be a continuation of the Neolithic Funnel Beaker culture), reinforced by recent genetic evidences showing shared European Neolithic farmer ancestry between both the cultures (Sintashta and Corded Ware) (*Allentoft et al., 2015*; *Wencel, 2015*), rationalize the finding of Funnel Beaker culture associated SORSUM sequence as the major recombination partner in RISE386 isolated from Sintashta remains. Besides, finding of ancestral genotype D sequence (like DA51 from the Saka) as minor recombination partner in aHBV from Sintashta people might be indicative of further eastward expansion of Sintashta people and admixture with the ancestors of the Saka people that perhaps inhibited the central steppe/central Asia (present day Kazakhstan/Kyrgyzstan) and carried the ancestral HBV that subsequently evolved into genotype D. It is important to note that majority of the aHBV sequences related to diverse subgenotypes of D (DA51, DA27, DA222, DA29) were isolated from different parts of central steppe/central Asia (**Table 1**), emphasizing the significance of this geographical area in origin, evolution and diversification of genotype D.

Another important finding of the present recombination analysis was the detection of DA51 (ancestral genotype D) and RISE386 (ancestral genotype A) like sequences within the extinct genotype sequences (RISE563 and RISE 254) exclusively from Bronze age (event RC, Table 2, Figure 3), but not in phylogenetically related sequences from preceding Neolithic Age (KARSDORF, SORSUM). This points toward probable early Bronze age epidemiological interaction between European people (then carrying the extinct HBV genotype) with people from Eurasian Steppe and Central Asian (carrying ancestral genotypes A & D). Thus, it appears that at least 3 distinct HBV genetic types would be existing concurrently during Bronze age - the Neolithic genotype (distributed in eastern & western Europe), primeval genotype A and D in different regions of Steppe/Central Asia. However, subsequent to Bronze age, the Neolithic genotype disappeared to be never found again, instead, ancestral genotype A sequences (DA195 and DA119) emerged in eastern European Iron age Hungarian Scythian and North Carpathian populations, respectively (Figure 5). Based on the absence of 6 nucleotide insertion at the carboxy terminal end of the core gene ORF in aHBV isolates RISE386, RISE387 and DA195, while its presence in isolate DA119 and in extant A sequences, DA119 has been proposed as the ancestor for all the modern subgenotypes of genotype A (*Mühlemann et al., 2018*). These observations evidently indicate a transition of HBV genotypes in European populations from Bronze age to Iron age, leading to the emergence of genotype A (from Steppe), a subgenotype (Ae) of which is presently predominant in Europe (*Kramvis, 2014*). This transition corroborates well with massive Bronze age ‘*Steppe migration*’ resulting in replacement of the Neolithic farming cultures of Europe, evident by recent genetic data demonstrating rapid replacement of the earlier European Y chromosome haplogroups by people from the east carrying *R1a* and *R1b* haplogroups, the latter being the most common haplogroups in present-day European populations (*Haak et al., 2015; Allentoft et al., 2015*). Nevertheless, despite extinction, remnants of the Neolithic HBV genotype do continue to exist as recombinant fragments in extant HBV genotypes (A, C, D, I, and E etc.), evocative of ancient origin of these genotypes.

**Figure 5.**
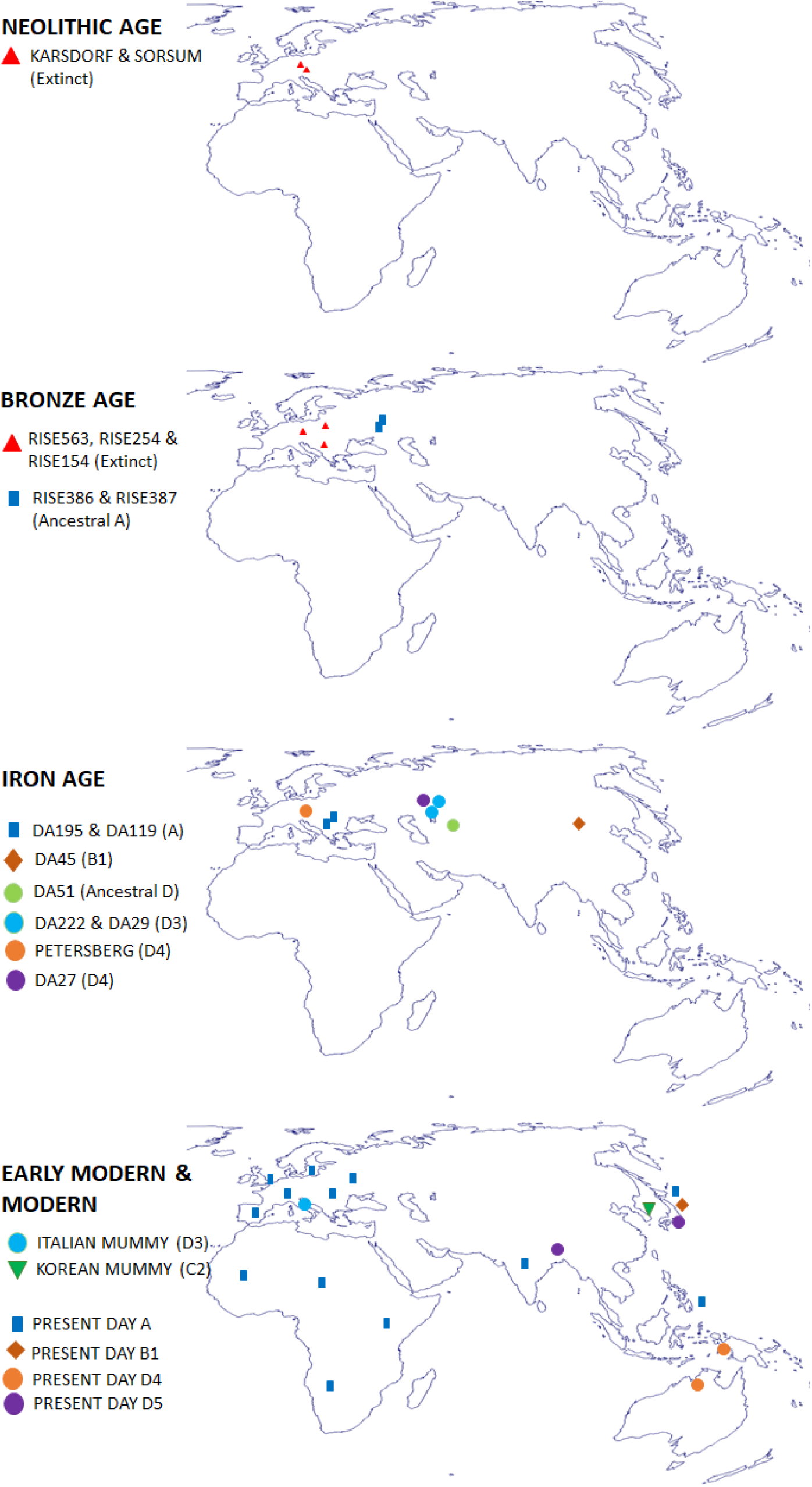
Maps showing the distribution of different HBV genotypes at different time periods, based on the aHBV record, available till now. The figure shows the expansion of the Neolithic genotype (presently extinct, denoted by red triangle) from Neolithic to Bronze age (upper two panels). Lower three panels show the migration of genotype A into Europe and subsequent wide expansion to current distribution. The lower two panels depict the Iron age distribution of certain subgenotypes of genotypes B (B1) and D (D4 & D5), as well as their presently restricted distribution. Base maps were https://d-maps.com/carte.php?num_car=3267&lang=en and annotated using Microsoft PowerPoint application.

On the other hand, analysis of the available aHBV genotype D sequences (DA51, DA27, DA222, PETERSBERG, DA29 and Italian mummy) revealed no significant shared recombination event(s) within these genomes, suggesting their independent evolution after divergence from the ancestral sequence. Analysis showing non-recombinant DA51 in nomadic Indo-Iranian Saka people from Kyrgyzstan, Central Asia at the basal position of extant genotype D clade and presence of at least 2 diverse subgenotypes of D namely D3 (DA29, DA222) and D5 (DA27) within the Iron age populations of central Asia (particularly Kazakhstan), raise doubts on the recently proposed North African/Middle Eastern origin of genotype D (*Kostaki et al., 2018*). From the available aHBV sequences, it seems more logical to reconsider central Asia as the site of origin of HBV genotype D in certain nomadic/warrior populations, followed by evolution of different subgenotypes and their dispersal with rapid expansion/invasion by these populations. Although, chronological appearance of subgenotype D5 precedes subgenotype D4 in the presently available aHBV sequence records, subgenotype D4 is believed to be the ancient one, associated with early intercontinental migrations of human (*Kramvis, 2014*). Nevertheless, distribution of both these subgenotypes in ancient population is discrepant with their current distributions as discussed earlier. At present, subgenotype D4 is restricted in geographically distant and isolated aboriginal populations of Australian, Micronesia, Papua New Guinea and Arctic Denes, which attests to its ancient origin (*Kramvis, 2014*). It has also been sporadically detected in some parts of Africa, and the Caribbean islands, which has been suggested to indicate either its ancient African origin or relatively recent introduction through slave trade (*Kramvis, 2014; Banerjee et al. 2014*). Recently, a regional variant of this subgenotype has also been reported from north eastern India (*Banerjee et al. 2014*), which might be a spill-over of the original subgenotype D4 *en route* pacific islands. However, with the present distribution of this subgenotype, its presence in Iron Age Germany (PETERSBERG) remains inexplicable. Similarly, more data is required to rationalize the Iron Age detection of subgenotypes B1 (DA45 from Mongolia), and D5 (DA27 from Kazakhstan) with their presently restricted distribution in Japan and Eastern India, respectively (*Kramvis, 2014; Banerjee et al., 2006; Ghosh et al., 2010*), although links to East Asian ancestry has been suggested to explain the phylogenetic similarities between isolate DA27 and subgenotype D5 sequences found among the Paharia people of Eastern India (*Muhlemann et al., 2018; Damgaard et al., 2018; Basu et al., 2016*). Nevertheless, these observations contradict classical belief that genotypes and subgenotypes have evolved in geographical regions or populations, presently found.

Among the most disagreed topics in HBV molecular evolution is its age estimate and over the years different research groups have proposed diverse models suggesting extensively different dates of origin, ranging from few hundred to thousand years (*Muhlemann et al., 2018; Krause-Kyora et al., 2018; Dominguez Souza et al., 2018; Zehender, 2015; Zehender et al., 2012*; *Jazayeri, 2010; Zhou & Holmes, 2007; Hannoun et al., 2005; Fares & Holmes, 2002; Bollyky & Holmes, 1999*). Nevertheless, most of these estimates have been reported to be seriously biased and too improbable to explain the present-day prevalence and geographical distribution of HBV (*Littlejohn et al., 2016*). Discoveries of aHBV sequences from Neolithic (*Krause-Kyora et al., 2018*) and Bronze Age (*Muhlemann et al., 2018*) have provided direct evidences to straightway reject a number of these theories (*Dominguez Souza et al., 2018; Zehender, 2015; Zehender et al., 2012*; *Jazayeri, 2010; Hannoun et al., 2005; Bollyky & Holmes, 1999*). Yet, time estimates for origin and divergence of HBV genotypes in *Muhlemann et al.*, also appear to be predisposed to serious underestimation. Such an underestimation is predictable, as time estimates reported in *Muhlemann et al.*, was based on analyses of viral phylogenies calibrated on heterochronous sample derived molecular clock estimates. Notably, *Muhlemann et al*., using root-to-tip regression and date randomization tests, reportedly found clear positive temporal signal in their dataset, which is imperative for an informative tip-dated analysis and to ascertain the suitability of molecular clock model for a given dataset. However, in sharp contrast, using same statistical tests, *Krause-Kyora et al., 2018* and *Ross-patterson et al., 2018* failed to detect satisfactory evidence of temporal signal in their datasets (with or without aHBV sequences), which questions the relevance of molecular clock models for time-scale analyses of HBV datasets. Inadequacy of temporal signal in HBV datasets is well anticipated, since evolutionary history of HBV involves horizontal gene-flow, widespread recombinations, point mutation, gene overprinting, as well as cross-species transmission (*Lauber et al., 2017; Krause-Kyora et al., 2018*). Since molecular clock based coalescent approaches represent short-term evolutionary rates and principally recommended for analysing short-term evolution of rapidly evolving viruses, this method has been described to be inappropriate for estimating the time of origin and divergence of HBV and other viruses having extremely deep coevolutionary history with their hosts (*Lauber et al., 2017; Ho et al., 2011; Sharp & Simmonds, 2011; Rambaut et al., 2016*). Use of molecular clock calibrated phylogenies have therefore been held responsible for highly conflicting date estimates obtained from analyses of extant as well as aHBV sequences (*Bouckaert et al., 2013; Pimenoff et al., 2018*).

In the light of available literature, divergence times estimated by *Muhlemann and colleagues* appear to be incompatible with diverse ancient HBV genotypes/subgenotypes that are believed to have emerged, co-evolved and dispersed ‘*Out of Africa*’ with human several thousand years ago (*Dominguez Souza et al., 2018; Littlejohn et al., 2016; Locarnini et al., 2013*). Most convincing and direct evidences emanates from the prevalence of distinct subgenotypes of genotype C in the isolated ‘*relict populations*’ (exclusive prevalence of C4 in Australian aborigines, C1 in *Jarawas* and *Orang Asli* from Andaman Island and Malaysia, respectively, C3 in Melanesians and C-14 like subgenotype in Torres Strait Islanders), who are believed to be the descendants of indigenous east Africans that followed the ‘*coastal trail*’ and populated these islands as early as 50,000-60,000 years ago or even earlier (*Yuen et al., 2019; Tobler et al., 2017; Littlejohn et al., 2016; Locarnini et al., 2013; Macaulay-Science-2005*). Notably, *Muhlemann* also could not find support for the divergence of subgenotype C3 and Haitian subgenotype A3 strains from their analyses, which indicates the incompatibility of the divergence estimates with the distribution of these subgenotypes. While rates estimated by *Muhlemann and colleagues* essentially signify short term evolution, for several viruses including hepadnaviruses, it is now well established that short-term rates vary extensively from longterm evolutionary and therefore, use of short term evolutionary rates are expected to result in serious underestimation of time-scale of origin, emergence, and dispersal, often differing by several orders of magnitude (*Duchêne et al., 2014; Lauber et al., 2017; Littlejohn et al., 2016; Zehender et al., 2014; Godoy et al., 2013; Feschotte_Gilbert, 2012; Sharp and Simmonds, 2011; Gilbert and Feschotte, 2010*). For that reason, HBV divergence estimated by *Muhlemann and colleagues* are probably underestimated by 3-4 orders of magnitude, since the range of evolutionary rate calculated by their approach ranges from 8.04×10^−6^ to 1.51×10^-5^ nucleotide substitutions per site per year, which varies widely from the long term evolutionary rates of HBV, ranging from 10^-8^ to 10^-9^ nucleotide substitutions per site per year, as estimated in previous studies (*Lauber et al., 2017; Littlejohn et al., 2016; Zehender et al., 2014; Feschotte_Gilbert, 2012; Sharp and Simmonds, 2011; Gilbert and Feschotte, 2010; Simmonds, 2001*). Thus, to obtain reliable time estimates for deep divergence events, calibrations of viral phylogenies on long-term evolutionary rates (*Zehender et al., 2014; Godoy et al, 2013; Sharp & Simmonds, 2011*) or on host co-divergence events has been strongly recommended (*Pimenoff et al., 2018*).

Recent discoveries of HBV related viruses from diverse classes of vertebrates, endogenous HBV (eHBV) elements and hepadnavirus-like retroelement within insect genomes (HEART1 and HEART2) have established an extremely deep rooted history of these viruses, dating back to several hundred million years ago (mya) (*Gong & Han, 2018; Dominguez Souza et al., 2018; Lauber et al., 2017; Dill et al., 2016; Suh et al., 2014; Gilbert and Feschotte, 2010*). A recent study, based on analyses of HBV phylogeny of conserved domains of HBV polymerase protein and molecular clock calibrated on endogenous Avihepadnaviral sequences, has demonstrated that the ancestral Orthohepadnaviruses (hepadnaviruses infecting mammalian hosts) diverged from its nearest clade of ancestral Metahepadnaviruses (hepadnaviruses infecting piscine hosts) approximately 235 mya, during the *Triassic* period (*Lauber et al., 2017*). Among the Orthohepadnaviruses, the New World HBV diverged from the ancestor of Old-World HBV around 55 mya, while the later started diverging into different human/NHP genotypes around 30 mya, subsequently co-evolving with diverse hosts belonging to the superfamily *Hominoidea (Lauber et al., 2017*). A subsequent study demonstrated natural infection of hepadnaviruses among capuchin monkeys (platyrrhine, New-World primate), and based on the recent findings of platyrrhine fossils records from Amazonian Peru (*Bond et al., 2015*) suggested that ancestral New World hepadnaviruses, phylogenetically related to Old-World human/NHP associated hepadnaviruses were introduced into the South America, probably with the trans-Atlantic migration of African pre-platyrrhines, at least 36 mya (*Dominguez Souza et al., 2018; Houle et al., 1999*). Similar deep co-evolutionary history of other dsDNA viruses with their hosts over several million years have been recognized in the recent years (*Pimenoff-Bravo, 2019; Pimenoff, 2018; Wertheim, 2014; Kolb et al., 2013*).

Till date, several studies have focussed on the origin, dispersal and geographic distribution of human HBV genotypes predominant in the old world (*Littlejohn et al., 2016; Rasche et al., 2016; Zehender et al., 2014*). On the contrary, dispersal of human genotypes F and H to the New World as well as dispersal of NHP HBV from Africa (considering the origin in the African continent) (*Rasche et al., 2016; Souza et al., 2014*) to distant parts of South East Asia, transatlantic South America remains debated (*Littlejohn et al., 2016; Souza et al., 2014*). It was earlier assumed that virus(s) ancestral to genotypes F and H might have travelled with human along their pre-historic migration to the Americas through the Bering Strait (*Zehender et al., 2014; Paraskevis et al., 2013*). But wide phylogenetic distance and absence of a related or common HBV strain (s) on both side of the *Beringia* do not seem to support this mode of intercontinental transmission of HBV (*Rasche et al., 2016*). On the other hand, despite being distributed in distantly located geographical areas, in the present phylogenetic network, clades of NHP HBV sequence from Africa, South East Asia and Africa were found to originate from a close common basal net, signifying deep shared ancestry among the NHP HBVs, while upholding considerable distance from Old world human HBV genotypes (Figure 1), supporting previous finding (*Hu et al., 2001*). Furthermore, in this analysis, New World primate HBV sequences (WMHBV and CMHBV) emerge as an offshoot from the clade representing human genotypes F and H (Figure 1). Phylogenetic relatedness of the genotypes F and H sequences with highly divergent WMHBV and CMHBV probably indicates introduction of a common or a group of related ancestral viruses into the New World, rather than a recent acquisition of genotypes F and H, distinct from WMHBV and CMHBV. Previous phylogenetic analyses, have also suggested an early divergence of genotype F (*Norder et al., 1996*) and WMHBV (*Bonvicino et al., 2014*) from rest of the human/NHP genotypes, indicating their ancient introduction in the New World (*Norder et al., 1996; Bailey et al., 1992*). Although, such an intercontinental introduction of HBV has been proposed to be caused by the transatlantic migration of HBV infected pre-platyrrhine primates (*Dominguez Souza et al., 2018*), but the requirement of specialized rafting skills and pre-adaptation to withstand extreme geographical and physiological conditions during long transoceanic migrations (*Bond et al., 2015; Houle et al., 1999*) raise serious doubts on the practicability of such proposals (*Heads, 2010; Simons, 1976*; *Fooden, 1972*).

An alternative to the transoceanic migration of the primates, Michael Heads (2010) proposed a model based on primate molecular phylogenies calibrated with radiometrically dated tectonic events. This model argues that the primate ancestral complex (early *Archonta*) was widespread across the supercontinent *Pangaea*, which differentiated with the break-up of *Pangaea* (∼185 mya) and subsequent geological events resulting in divergence, evolution, and present-day distribution of modern primate clades (*Heads, 2010*). This model is also supported by previous findings from extensive analyses of vertebrate molecular data, which suggested that majority of the modern mammalian orders had already diversified before the Cretaceous/Tertiary extinction and that continental break-up during the Cretaceous played an important role in diversification and distribution of the modern mammalian orders (*Hedges, 2002; Kumar & Hedges, 1998; Springer et al., 1997; Hedges et al., 1996; Fooden, 1972*). Considering the estimated Triassic origin of mammalian hepadnaviruses (*Orthohepadnavirus*) approximately 235 mya (*Lauber et al., 2017*), long before the break-up of Pangaea (*McIntyre et al., 2017; Heads, 2010*), it is tempting to assume that the ancestral Orthohepadnaviruses might have infected the primate ancestors widespread across Pangaea, which with subsequent break-up of Pangaea and other geologic events, started diverging and evolving independently in different geographic areas along with their hosts. Unlike previous models, this model can explain the present-day distribution of phylogenetically closely related hepadnaviruses among primates in distant continents, without necessitating to assume that their ancestors undertook impracticably long-distance intercontinental migrations or trans-oceanic voyages. Likewise, different viruses under the family *Herpesviridae* have also been proposed to have diverged with the split of Pangaea and subsequent coevolution in different primate groups (*Grose, 2012*).

Consequent to an earlier proposal (*Littlejohn et al., 2016*), recent findings strongly imply that members in the genus *Homo*, including *Neanderthals, Denisovans, Homo floresiensis (Brown et al., 2004)*, and the newly described *Homo luzonensis* (*Detroit et al., 2019*) etc. might have also carried archaic HBV genotypes, conceivably distinct from those carried by the early anatomically modern human (AMH) and the extant genotypes, similar to that suggested for papillomaviruses and herpesviruses (*Pimenoff et al., 2018*). Furthermore, considering the proportion of DNA introgressed from *Neanderthals* and *Denisovans* into modern humans, it remains important to review, if HBV genotypes presently circulating in Europe originated from Neanderthals, as speculated for papillomaviruses and herpesviruses (*Pimenoff et al., 2018*), while those circulating in islands east of Wallace’s Line originated from Denisovans or perhaps from any of the recently discovered hominins from SE Asia (*Detroit et al., 2019; Yuan et al., 2017; Cooper and Stringer, 2013; Brown et al., 2004*). Archaeologic evidences have clearly established that these archaic human species have co-existed during the time when AMH were expanding and peopling different regions, including the regions where remains from these archaic human species were found (*Detroit et al., 2019; Sutikna et al., 2016; Reich et al., 2010; Green et al., 2010*). Additionally, genetic studies have now proven that among the archaic human species, at least Neanderthals and Denisovans interbred among themselves as well as with the AMH, resulting in introgression of genetic material, and possibly the sexually transmitted viruses (*Pimenoff, 2018; Slon et al., 2018; Wolf and Akey, 2018; Villanea and Schraiber, 2018; Chen et al., 2017*; *Sankararaman et al., 2016; Sankararaman et al., 2014; Cooper and Stringer, 2013; Locarnini et al., 2013*).

In conclusion, ancient pathogen genomics has unbolted the doors to understanding origin, divergence and distribution of pathogens including HBV. Therefore, it will be important to search for aHBV genomes in existing archaic human/ hominin whole genome trace files and in yet to be sequenced DNA samples from archaic human remains to better understand the distribution and dispersal of HBV through time scales. The present research demonstrates that systematic analyses of aHBV sequences and interpretation of the findings from the context of relevant human genetic data can reveal interesting facts not only about origin, divergence and distribution of ancient pathogens, but can also provide clues about ancient human migration patterns.

## Acknowledgements

The author expresses sincere gratitude to Prof. Anna Kramvis, University of Witwatersrand, South Africa, Dr. Barbara Muhlemann, University of Cambridge, United Kingdom and Prof. Ben Krause-Kyora, Kiel University, Germany for generously sharing HBV DNA datasets. The author thankfully acknowledges Dr. Sonika Sharma, Defence Research Laboratory, Tezpur, India for critical review of the manuscript and constructive discussions.

## Data availability

Dataset used for reconstruction of phylogenetic network and recombination analyses, recombination result file (RDP4 output file) are available from the author upon request.

## References

Allentoft, M.E. et al. Population genomics of Bronze Age Eurasia. Nature 522, 167–172 (2015).

Arankalle, V.A. et al. A novel HBV recombinant (genotype I) similar to Vietnam/Laos in a primitive tribe in eastern India. J. Viral. Hepat. 17, 501–510 (2010).

Araujo, N.M. Hepatitis B virus intergenotypic recombinants worldwide: An overview. Infect. Genet. Evol. 36, 500–510 (2015).

Bailey, W.J. et al. Reexamination of the African hominoid trichotomy with additional sequences from the primate beta-globin gene cluster. Mol. Phylogenet. Evol. 1, 97–135 (1992).

Banerjee, P. et al. A rare HBV subgenotype D4 with unique genomic signatures identified in North-Eastern India –An emerging clinical challenge? PLoS One 9, e109425 (2014).

Bar-Gal, G.K. et al. Tracing hepatitis B virus to the 16th century in a Korean mummy. Hepatology 56, 1671–1680 (2012).

Basu, A., Sarkar-Roy, N. & Majumder, P. P. Genomic reconstruction of the history of extant populations of India reveals five distinct ancestral components and a complex structure. Proc. Natl Acad. Sci. USA 113, 1594–1599 (2016).

Bellwood, P. The search for ancient DNA heads east. Science 361, 31–32 (2018).

Bollyky, P.L & Holmes, E.C. Reconstructing the complex evolutionary history of hepatitis B virus. J. Mol. Evol. 49, 130–41 (1999).

Bond, M. et al. Eocene primates of South America and the African origins of New World monkeys. Nature 520, 538–541 (2015).

Bonvicino, C.R., Miguel, A.M. & Soares, M.A. Hepatitis B virus lineages in mammalian hosts: Potential for bidirectional cross-species transmission. World. J. Gastroenterol. 20, 7665–7674 (2014).

Bouckaert, R., Alvarado-Mora, M.V. & Pinho, J.R. Evolutionary rates and HBV: issues of rate estimation with Bayesian molecular methods. Antivir. Ther. 18, 497–503 (2013).

Brown, P. et al. A new small-bodied hominin from the Late Pleistocene of Flores, Indonesia. Nature 431, 1055–1061 (2004).

Chen, Z. et al. Ancient evolution and dispersion of human Papillomavirus 58 variants. J. Virol. 91, e01285–17 (2017).

Cooper, A. & Stringer, C.B. Did the Denisovans cross Wallace’s Line? Science 342, 321–323 (2013).

Damgaard, P.B. et al. 137 ancient human genomes from across the Eurasian steppes. Nature 557, 369–374 (2018).

Détroit, F. et al. A new species of Homo from the Late Pleistocene of the Philippines. Nature 568, 181–186 (2019).

Dill, J.A. et al. Distinct viral lineages from fish and amphibians reveal the complex evolutionary history of hepadnaviruses. J. Virol. 90, 7920–7933 (2016).

Dominguez Souza, B.F. de Carvalho. et al. A novel hepatitis B virus species discovered in capuchin monkeys sheds new light on the evolution of primate hepadnaviruses. J. Hepatol. 68, 1114–1122 (2018).

Duchêne, S., Holmes, E. C. & Ho, S. Y.W. Analyses of evolutionary dynamics in viruses are hindered by a time-dependent bias in rate estimates. Proc. R. Soc. Lond. B 281, 20140732 (2014).

Fang, Z.L. et al. A complex hepatitis B virus (X/C) recombinant is common in Long An county, Guangxi and may have originated in southern China. J. Gen. Virol. 92, 402–411 (2011).

Fares, M.A. & Holmes, E.C. A revised evolutionary history of hepatitis B virus (HBV). J. Mol. Evol. 54, 807–814 (2002).

Feschotte, C. & Gilbert, C. Endogenous viruses: insights into viral evolution and impact on host biology. Nature Rev. Genet. 13, 283–296 (2012).

Fooden, J. Breakup of Pangaea and isolation of relict mammals in Australia, South America, and Madagascar. Science 175, 894–898 (1972).

Ghosh, S. et al. Unique hepatitis B virus subgenotype in a primitive tribal community in eastern India. J. Clin. Microbiol. 48, 4063–4071 (2010).

Gilbert, C. & Feschotte, C. Genomic fossils calibrate the long-term evolution of hepadnaviruses. PLoS Biol. 8, e1000495 (2010).

Godoy, B.A., Alvarado-Mora, M.V., Gomes-Gouvêa, M.S., Pinho, J.R. & Fagundes, N. Origin of HBV and its arrival in the Americas-the importance of natural selection on time estimates. Antivir. Ther. 18, 505–512 (2013).

Gong, Z. & Han, G.Z. Insect retroelements provide novel insights into the origin of hepatitis B viruses. Mol. Biol. Evol. https://doi:10.1093/molbev/msy129 (2018).

Green, R.E. et al. A draft sequence of the Neandertal genome. Science 328, 710–722 (2010).

Grose, C. Pangaea and the Out-of-Africa Model of Varicella-Zoster Virus Evolution and Phylogeography. J. Virol. 86, 9558–9565 (2012).

Haak, W. et al. Massive migration from the steppe was a source for Indo-European languages in Europe. Nature 522, 207–211 (2015).

Haldipur, B.P., Walimbe, A.M. & Arankalle, V.A. Circulation of genotype-I hepatitis B virus in the primitive tribes of Arunachal Pradesh in early sixties and molecular evolution of genotype-I. Infect. Genet. Evol. 27, 366–374 (2014).

Hall, T.A. BioEdit: a user-friendly biological sequence alignment editor and analysis program for Windows 95/98/NT. Nucl. Acids. Symp. Ser. 41, 95–98 (1999).

Hannoun, C., Norder, H. & Lindh, M. An aberrant genotype revealed in recombinant hepatitis B virus strains from Vietnam. J. Gen. Virol. 81, 2267–2272 (2000).

Hannoun, C., Söderström, A., Norkrans, G. & Lindh, M. Phylogeny of African complete genomes reveals a West African genotype A subtype of hepatitis B virus and relatedness between Somali and Asian A1 sequences. J. Gen. Virol. 86, 2163–2167 (2005).

Heads, M. Evolution and biogeography of primates: a new model based on molecular phylogenetics, vicariance and plate tectonics. Zoologica Scripta 39, 107–127 (2010).

Hedges, S.B. The origin and evolution of model organisms. Nature Rev. Genet. 3, 838–849. (2002).

Hedges, S.B., Parker, P.H., Sibley, C.G. & Kumar, S. Continental breakup and the ordinal diversification of birds and mammals. Nature 381, 226–229 (1996).

Ho, S.Y. et al. Time-dependent rates of molecular evolution. Mol Ecol. 20, 3087–3101 (2011).

Houle, A. The origin of platyrrhines: An evaluation of the Antarctic scenario and the floating island model. Am. J. Phys. Anthropol. 109, 541–559 (1999).

Hu, X., Javadian, A., Gagneux, P. & Robertson, B.H. Paired chimpanzee hepatitis B virus (ChHBV) and mtDNA sequences suggest different ChHBV genetic variants are found in geographically distinct chimpanzee subspecies. Virus Res. 79, 103–108 (2001).

Huson, D.H. & Bryant, D. Application of phylogenetic networks in evolutionary studies. Mol. Biol. Evol. 23, 254–267 (2006).

Jazayeri, S.M., Alavian, S.M. &, Carman, W.F. Hepatitis B virus: origin and evolution. J. Viral. Hepat. 17, 229–235 (2010).

Kolb, A.W. Ane, C. & Brandt, C.R. Using HSV-1 genome phylogenetics to track past human migrations. PLoS One 8, e76267 (2013).

Kostaki, E.G. et al. Unravelling the history of hepatitis B virus genotypes A and D infection using a full-genome phylogenetic and phylogeographic approach. eLife 7, e36709 (2018).

Kramvis, A. Genotypes and Genetic Variability of Hepatitis B Virus. Intervirology 57, 141–150 (2014).

Krause-Kyora, B. et al. Neolithic and Medieval virus genomes reveal complex evolution of hepatitis B. eLife 7, e36666 (2018).

Kumar, S. & Hedges, S.B. A molecular timescale for vertebrate evolution. Nature 392, 917–920 (1998).

Lauber, C. et al. Deciphering the origin and evolution of Hepatitis B Viruses by means of a family of non-enveloped fish viruses. Cell. Host. Microbe. 22, 387–399 (2017).

Littlejohn, M., Locarnini, S. & Yuen, L. Origins and evolution of Hepatitis B Virus and Hepatitis D Virus. Cold. Spring. Harb. Perspect. Med. 6, a021360 (2016).

Locarnini, S., Littlejohn, M., Aziz, M.N. & Yuen, L. Possible origins and evolution of the hepatitis B virus (HBV). Semin. Cancer. Biol. 23, 561–575 (2013).

Macaulay, V. et al. Single, rapid coastal settlement of Asia revealed by analysis of complete mitochondrial genomes. Science 308, 1034–1036 (2005).

Magiorkinis, E.N., Magiorkinis, G.N., Paraskevis, D.N. & Hatzakis, A.E. Re-analysis of a human hepatitis B virus (HBV) isolate from an East African wild born *Pan troglodytes schweinfurthii*: evidence for interspecies recombination between HBV infecting chimpanzee and human. Gene 349, 165–171 (2005).

Makarenkov, V. & Legendre, P. From a phylogenetic tree to a reticulated network. J. Comput. Biol. 11, 195–212 (2004).

Martin, D.P., Murrell, B., Golden, M., Khoosal, A. & Muhire, B. RDP4: Detection and analysis of recombination patterns in virus genomes. Virus Evol. 1, vev003 (2015).

McIntyre, S.R.N., Lineweaver, C.H., Groves, C.P. & Chopra, A. Global biogeography since Pangaea. Proc. Biol. Sci. 284, 20170716 (2017).

Mühlemann, B. et al. Ancient hepatitis B viruses from the Bronze age to the Medieval period. Nature 557, 418–423 (2018).

Norder, H., Ebert, J.W., Fields, H.A., Mushahwar, I.K. & Magnius, L.O. Complete sequencing of a gibbon hepatitis B virus genome reveals a unique genotype distantly related to the chimpanzee hepatitis B virus. Virology 218, 214–223 (1996).

Olinger, C.M. et al. Possible new hepatitis B virus genotype, southeast Asia. Emerg. Infect. Dis. 14, 1777–1780 (2008).

Paraskevis, D. et al. Dating the origin and dispersal of hepatitis B virus infection in humans and primates. Hepatology 57, 908–916 (2013).

Pimenoff, V.N. & Bravo, I.G. Sexual Transmission of HPVs from Neanderthals to Modern Humans. HPV World 77 http://www.hpvworld.com/media/29/media_section/8/0/1080/pimenoff.pdf (2019).

Pimenoff, V.N., Houldcroft, C.J., Rifkin, R.F. & Underdown, S. The Role of aDNA in Understanding the Coevolutionary Patterns of Human Sexually Transmitted Infections. Genes (Basel) 9, E317 (2018).

Pourkarim, M.R., Amini-Bavil-Olyaee, S., Kurbanov, F., Van Ranst, M. & Tacke, F. Molecular identification of hepatitis B virus genotypes/subgenotypes: revised classification hurdles and updated resolutions. World. J. Gastroenterol. 20, 7152–7168 (2014).

Rambaut, A., Lam, T.T., Max Carvalho, L. & Pybus, O.G. Exploring the temporal structure of heterochronous sequences using TempEst (formerly Path-O-Gen). Virus Evol. 2, vew007 (2016).

Rasche, A., Souza, B.F.C.D. & Drexler, J.F. Bat hepadnaviruses and the origins of primate hepatitis B viruses. Curr. Opin. Virol. 16, 86–94 (2016).

Reich, D. et al. Genetic history of an archaic hominin group from Denisova Cave in Siberia. Nature 468, 1053–1060 (2010).

Ross, Z.P. et al. The paradox of HBV evolution as revealed from a 16th century mummy. PLoS Pathog. 14, e1006750 (2018).

Sankararaman, S. et al. The genomic landscape of Neanderthal ancestry in present-day humans. Nature 507, 354–357 (2014).

Sankararaman, S., Mallick, S., Patterson, N. & Reich, D. The combined landscape of Denisovan and Neanderthal ancestry in present-day humans. Curr. Biol. 26, 1241–1247 (2016).

Sharp, P.M. & Simmonds, P. Evaluating the evidence for virus/host co-evolution. Curr. Opin. Virol. 1, 436–41 (2011).

Shi, W. et al. Subgenotyping of genotype C Hepatitis B Virus: correcting misclassifications and identifying a novel subgenotype. PLoS One 7, e47271 (2012).

Simmonds, P. & Midgley, S. Recombination in the genesis and evolution of hepatitis B virus genotypes. J. Virol. 79, 15467–15476 (2005).

Simmonds, P. The origin and evolution of hepatitis viruses in humans. J. Gen. Virol. 82, 693–712 (2001).

Simons, E.L. The fossil record of primate phylogeny. Molecular Anthropology (Springer Press, 1976).

Slon, V. et al. The genome of the offspring of a Neanderthal mother and a Denisovan father. Nature 561, 113–116 (2018).

Souza, B.F., Drexler, J.F., Lima, R.S., Rosario, M.O. & Netto, E.M. Theories about evolutionary origins of human hepatitis B virus in primates and humans. Braz. J. Infect. Dis. 18, 535–543 (2014).

Springer, M.S. et al. Endemic African mammals shake the phylogenetic tree. Nature 388, 61–64 (1997).

Spyrou, M.A., Bos, K.I., Herbig, A. & Krause, J. Ancient pathogen genomics as an emerging tool for infectious disease research. Nature Rev. Genet. 20, 323–340 (2019).

Su, H. et al. A novel complex A/C/G intergenotypic recombinant of hepatitis B virus isolated in southern China. PLoS One 9, e84005 (2014).

Suh, A. et al. Early mesozoic coexistence of amniotes and hepadnaviridae. PLoS Genet. 10, e1004559 (2014).

Sutikna, T. et al. Revised stratigraphy and chronology for Homo floresiensis at Liang Bua in Indonesia. Nature 532, 366–369 (2016).

Tamura, K., Stecher, G., Peterson, D., Filipski, A. & Kumar, S. MEGA6: Molecular Evolutionary Genetics Analysis version 6.0. Mol. Biol. Evol. 30, 2725–2729 (2013).

Tobler, R. et al. Aboriginal mitogenomes reveal 50,000 years of regionalism in Australia. Nature 544, 180–184 (2017).

Tran, T.T., Trinh, T.N. & Abe, K. New complex recombinant genotype of hepatitis B virus identified in Vietnam. J. Virol. 82, 5657–5563 (2008).

Vartanian, J.P. et al. Identification of a hepatitis B virus genome in wild chimpanzees (*Pan troglodytes schweinfurthi*) from East Africa indicates a wide geographical dispersion among equatorial African primates. J. Virol. 76, 11155–11158 (2002).

Villanea, F.A. & Schraiber, J.G. Multiple episodes of interbreeding between Neanderthal and modern humans. Nature Ecol. Evol. 3, 39–44 (2019).

Wencel, M.M. An absolute chronological framework for the central-eastern European Eneolithic. Ox. J. Archaeol. 34, 33–43 (2015).

Wertheim, J.O. et al. Evolutionary origins of human herpes simplex viruses 1 and 2. Mol. Biol. Evol. 31, 2356–2364 (2014).

Wolf, A.B. & Akey, J.M. Outstanding questions in the study of archaic hominin admixture. PLoS Genet. 14, e1007349 (2018).

Yuan, D. et a. Modern human origins: multiregional evolution of autosomes and East Asia origin of Y and mtDNA. Preprint at https://www.biorxiv.org/content/10.1101/101410v5 (2018).

Yuen, L.K.W. et al. Tracing ancient human migrations into Sahul using Hepatitis B Virus genomes. Mol. Biol. Evol. 36, 942–954 (2019).

Zehender, G. et al. Enigmatic origin of hepatitis B virus: an ancient travelling companion or a recent encounter? World. J. Gastroenterol. 20, 7622–7634 (2014).

Zehender, G. et al. Reliable timescale inference of HBV genotype A origin and phylodynamics. Infect. Genet. Evol. 32, 361–369 (2015).

Zehender, G. et al. Spatial and temporal dynamics of hepatitis B virus D genotype in Europe and the Mediterranean Basin. PLoS One 7, e37198 (2012).

Zhou, Y. & Holmes, E.C. Bayesian estimates of the evolutionary rate and age of hepatitis B virus. J. Mol. Evol. 65, 197–205 (2007).

